# Seedling leaves allocate lower fractions of nitrogen to photosynthetic apparatus in nitrogen fixing trees (*Dalbergia odorifera and Erythrophleum fordii*) than in non-nitrogen fixing trees (*Betula alnoides and Castanopsis hystrix*) in subtropical China

**DOI:** 10.1101/482984

**Authors:** Jingchao Tang, Baodi Sun, Ruimei Cheng, Zuomin Shi, Da Luo, Shirong Liu, Mauro Centritto

## Abstract

Photosynthetic-nitrogen use efficiency (PNUE) is a useful trait to characterize leaf economics, physiology, and strategy. In this study, we investigated the differences in PNUE, leaf nitrogen (N) allocation, and mesophyll conductance (*g*_m_) in *Dalbergia odorifera* and *Erythrophleum fordii* (N-fixing trees), and *Betula alnoides* and *Castanopsis hystrix* (non-N-fixing trees). Seedlings of the four species were cultured in pots and received the same nutrient solution, water volume, and light. LiCor-6400 was used to determine fluorescence yield, photosynthetic response to light, and intercellular CO_2_ concentration (*C*_i_). N allocation fractions in the photosynthetic apparatus were calculated according to Niinemets and Tenhunen method; *g*_m_ was calculated according to variable *J*, EDO, and *A*-*C*_i_ curve fitting methods. PNUE of *D. odorifera* and *E. fordii* were significantly lower than those of *B. alnoides* and *C. hystrix* because of their allocation of a lower fraction of leaf N to Rubisco (*P*_R_) and bioenergetics (*P*_B_). Mesophyll conductance had a significant positive correlation with PNUE in *D. odorifera, E. fordii*, and *B. alnoides*. The fraction of leaf N to cell wall (*P*_CW_) had a significant negative correlation with *P*_R_ in *B. alnoides* and *C. hystrix*. We conclude that *B. alnoides* and *C. hystrix* optimized their leaf N allocation toward photosynthesis, with the trade-off being N allocation to the cell wall and Rubisco. Thus, these two species may have a higher competitive ability in natural ecosystems with fertile soil.

## Introduction

Nitrogen (N) is a major constituent of proteins, nucleic acids, amino acids, and chlorophylls [1,2] and is important for plant survival and growth. In leaves, N is mainly used for photosynthesis [3], and therefore, there is a clear relationship between leaf N contents and photosynthetic capacity. Many researchers use photosynthetic-N use efficiency (PNUE, the ratio of light-saturated net CO_2_ assimilation rate (*A*_max_′) to leaf N content per area (*N*_area_) [4]) to show how efficiently N resources are used during photosynthesis, and studies have been conducted on a variety of species [3,5,6].

N involved in photosynthesis has always been an important factor that influences PNUE [7]. Rubisco is the most abundant enzyme in C_3_ plants [8], and it is the key factor in carbon assimilation [9]. Many researchers have found a positive correlation between leaf N fraction in Rubisco (*P*_R_) and PNUE in various plants [10-12]. Apart from photosynthetic machinery, N in leaves is also involved in other machineries, such as structure, storage, and respiration [13]. Cell walls contain proteins that have functions in plant defense, growth, development, signaling, intercellular communication, environmental sensing, and as selective exchange interfaces [14, 15]. Trade-offs may occur between N allocation to cell walls, Rubisco (when N is needed to maintain cellular functions), seed development, and immune response [15-17]. However, some studies have shown that these trade-offs only exist in individuals of the same species [12] or species lacking N in leaves [17, 18].

CO_2_ is an important raw material for photosynthesis [19], and CO_2_ partial pressure is important for Rubisco activity; this is because O_2_ is a competitive inhibitor of the C assimilatory reaction of Rubisco, promoting the Rubisco oxidation reaction [20]. A significant negative correlation between *C*_i_ (intercellular CO_2_ concentration)-*C*_c_ (CO_2_ concentration at carboxylation site) and PNUE was found in *Populus cathayana* [21]. N is also involved in carbonic anhydrases and aquaporins [22]. These proteins play a role in mesophyll conductance (*g*_m_) by changing the nature of the diffusing molecule [23] and facilitating CO_2_ diffusion through membranes [24].Therefore, PNUE may be influenced by *g*_m_ [22]. A significant positive correlation was found between mesophyll conductance (*g*_m_) and PNUE in six *Populus* genotypes [25].

Species with a high *N*_area_ usually have a high *A*_max_′ according to the worldwide leaf economic spectrum [4]. However, there are some differences in N-fixing plants. N-fixing species could convert N from the air through legume bacteria, and this could meet more than half of the plant’s N demand. Several studies have shown that N-fixing species always have enough N in leaves [26-28]. One study showed that N-fixing trees had higher *A*_max_′ and *N*_area_, and there was no significant difference in PNUE or the proportion of leaf N allocated to Rubisco (*P*_R_) between N-fixing and non-N-fixing trees [29]. However, other studies have shown that N-fixing trees had lower *A*_max_′ and higher *N*_area_, which resulted in a lower PNUE [30, 31]. These contradicting results may imply that some N-fixing species use a different strategy to utilize N compared to non-N-fixing species. One possible explanation is that the percentage of N in the photosynthetic apparatus is lower in the N-fixing trees [30, 31]. However, these studies neglect that *g*_m_ and *P*_CW_ could also influence PNUE [15, 16, 32]. We studied the factors that affect PNUE in both N-fixing and non-N-fixing large trees and found *P*_R_ and *P*_B_ to be the main factors; the effects of *g*_m_ and *P*_CW_ were relatively small [33], but the effects in N-fixing tree seedlings remained unclear.

*Dalbergia odorifera*, *Erythrophleum fordii*, *Betula alnoides*, and *Castanopsis hystrix* are suitable for forestation in southern subtropical China and have high economic values [34-37]. *D. odorifera* and *E. fordii* are both evergreen N-fixing trees, whereas *B. alnoides* and *C. hystrix* are both non-N-fixing, and deciduous and evergreen, respectively. The objectives of our study are as follows: 1) understand how PNUE varies among *D. odorifera*, *E. fordii*, *B. alnoides*, and *C. hystrix* seedlings; 2) quantify the relationship between PNUE and photosynthetic capacity, leaf N allocation, and diffusional conductances to CO_2_, and 3) determine the relationship between N in Rubisco and the cell wall, and the relationship between PNUE and *g*_m_ in seedlings.

## Materials and Methods

### Study Area and Plant Material

This study was carried out in Experimental Center of Tropical Forestry (22°7′19″–22°7′22″N, 106°44′40″–106°44′44″E) of the Chinese Academy of Forestry located in Guangxi Pingxiang, China. The location has a subtropical monsoon climate with distinct dry and wet periods where the mean annual temperature is 21°C. The mean monthly minimum and maximum temperatures are 12.1°C and 26.3°C. The mean annual precipitation is 1400 mm, and it occurs mainly from April to September. Active accumulated temperature above 10°C is 6000–7600°C. The total annual sunshine duration is 1419 hours [38,39].

Seeds of *D. odorifera*, *E. fordii*, and *C. hystrix* were collected from a single tree for each species, and *B. alnoides* seedlings were somaclone. The seeds of *D. odorifera*, *E. fordii*, and *C. hystrix* were germinated in a seedbed in February 2014 and *B. alnoides* went through budding at the same time. When the seedlings were approximately 20 cm tall, 30 similarly sized seedlings per species were individually transplanted to pots (5.4 L, filled with washed river sand) and established in an open site at the Experimental Center of Tropical Forestry in March 2014. From April to June, each pot received the same nutrient solution (0.125 _g_ N and 0.11 _g_ P, Hyponex M. Scott & Sons, Marysville, OH, USA) once a week, and was watered every day to keep the soil moist. Natural light (100% of light in the field) was used for illumination.

### Determination of Gas Exchange Measurements

Gas exchange parameters were determined with a LiCor-6400 portable photosynthesis system (LI-COR, Lincoln, NE, USA) on sunny days from 8 am to 10 am in July and August 2014. Seven healthy and newly emerged leaves exposed to the sun in each tree species were chosen (one leaf per individual healthy tree). Photosynthetic response to photosynthetic photon flux density (PPFD) and intercellular CO_2_ concentration (*C*_i_, μmol mol–1) were determined for each leaf (seven repetitions in each species) according to Broeckx *et al.* 2014 [25] and Zhang *et al.* 2016 [40]. Relative humidity of the air in the leaf chamber was maintained at 60–70%, and leaf temperature was set at 30°C. The net photosynthetic rate (*A*_n_, μmol m^−2^ s^−1^), stomatal conductance (*g*_s_, mol CO_2_ m^−2^ s^−1^), and *C*_i_ of each sampled leaf were recorded ten times after 200 s under each PPFD and CO_2_ concentration. Then light-saturated net CO_2_ assimilation rate (*A*_max_′, μmol m^−2^ s^−1^), light-saturated day respiration rate (*R*_d_, μmol m^−2^ s^−1^) and light-and CO_2_-saturated net CO_2_ assimilation rate (*A*_max_, μmol m^−2^ s^−1^) were measured or calculated. For further details see Tang *et al*. [33].

### Determination of chlorophyll fluorescence and mesophyll conductance

Fluorescence yield was measured with a LiCor–6400 leaf chamber fluorometer (6400-40, LI-COR, Lincoln, Nebraska, USA), using the same leaf with seven repetitions of each species. Chamber temperature was maintained at 28–32°C, and chamber air relative humidity was maintained at 60–70%. Chamber CO_2_ concentration was set to 380 μmol mol^−1^. *PPFD* was set to light saturation point. Constant values of fluorescence yield (⊿*F*/*F*_m_′) of each leaf sample were recorded 10 times after 200 s [41]. We used Loreto *et al*. [42] methods to calculate the photosynthetic electron transport rate (*J*_f_, μmol m^−2^ s^−1^):

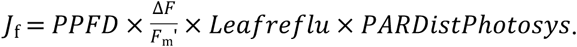

*Leafreflu* (leaf absorptance valued) and *PARDistPhotosys* (the fraction of quanta absorbed by photosystem II) were 0.85 [43] and 0.5 [42], respectively. We used the variable *J* method described by Harley *et al*. [44], which has been used in recent years [45-48] to calculate mesophyll conductance (*g*_m_, mol CO_2_ m^−2^ s^−1^):

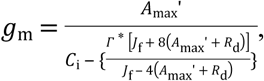

*R*_d_, *C*_i_, and *A*_max_*′* were determined from gas exchange measurements. The CO_2_ photo compensation point (*Γ*^*^,μmol mol^−1^) value was 54.76 at 30°C according to Bernacchi *et al* [49].

Because the Harley method should calibrate the ETR using Chl fluorescence and gas exchange under low O_2_, we used the experience value instead (*Leafreflu* = 0.85) [33]. We also used Ethier and Livingston [50] and the exhaustive dual optimization (EDO) method [51] to calculate *g*_m_. We used software based on the Ethier and Livingston method developed by Sharkey *et al.* [52] to get *g*_m_, and uploaded our data through a website (http://www.leafweb.org) to get *g*_m_ calculated by the EDO method.

### Determination of ***V*_cmax_ and *J*_max_**

The mean value of *g*_m_ calculated from three methods was used to calculate CO_2_ concentration in chloroplasts (*C*_c_, μmol mol^−1^):

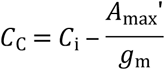

Then *C*_c_ was used to fit an *A*_n_-*C*_c_ curve, followed by the maximum carboxylation rate (*V*_cmax_, μmol m^−2^ s^−1^) calculated according to Farquhar *et al.* [9], and the maximum electron transport rate (*J*_max_, μmol m^−2^ s^−1^) calculated according to Loustau *et al.* [53]. The fitting model used *in vivo* Rubisco kinetics parameters (*K*_o_, *K*_c_, and their activation energy) measured by Niinemets and Tenhunen [7].

### Analysis of quantitative limitations of photosynthetic capacity

The relative controls on photosynthetic capacity imposed by stomatal conductance (*l*_s_, %), mesophyll diffusion (*l*_m_, %), and biochemical capacity (*l*_b_, %) were calculated following the quantitative limitation analysis of Grassi and Magnani [54] as applied in Tomás *et al* [55], Peguero-Pina *et al*. [56, 57] and Nha *et al*. [58]. Different fractional limitations, *l*_s_, *l*_m_, and *l*_b_ (*l*_s_ + *l*_m_ + *l*_b_ = 1) were calculated as:

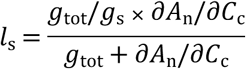

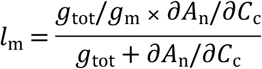

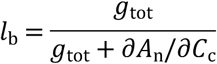

Where *g*_s_ and *g*_m_ were used in light-saturated and atmospheric CO_2_ concentration was 380 μmol mol^−1^, and *g*_m_ was the mean value of three methods. *g*_tot_ is the total conductance to CO_2_ from ambient air to chloroplasts (the sum of the inverse CO_2_ serial conductances *g*_s_ and *g*_m_). *∂A*_n_/*∂C*_c_ was calculated as the slope of *A*_n_–*C*_c_ response curves over a *C*_c_ range of 50–100 μmol mol^−1^ [56, 57].

### Determination of Additional Leaf Traits

Leaf samples used for gas exchange measurements and leaves which size was similar to leaves used for determine photosynthesis was taken. Leaf areas were measured with a scanner (Perfection v700 Photo, Epson, Nagano-ken, Japan). Leaf dry weights were measured using an analytic balance after being oven-dried at 80°C for 48 h, then leaf mass per area (LMA, g m^−2^) was calculated.

Dried leaf samples were ground into a dry flour. Organic carbon (C) concentration was determined by the potassium dichromate-sulfuric acid oxidation method (*C*_mass_ mg g^−1^). N concentration was determined by a VELP automatic Kjeldahl N determination apparatus (UDK-139, Milano, Italy), and leaf N per mass (*N*_mass_, mg g^−1^) and per area (*N*_area_ g·m^−2^) values were calculated [33]. PNUE (μmol·mol^−1^·s^−1^) was then calculated by the formula:

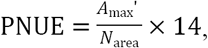

where 14 is the atomic mass of nitrogen.

Chlorophylls were extracted by direct immersion: 0.2 g of frozen leaves were cut into small pieces which were 5–10 mg. Leaf pieces were placed into a volumetric flask and 25 mL of 95% (v/v) alcohol was added. The flask was kept in the dark for 24 h. The absorbance of the extracts was measured at 665 nm and 649 nm with a Shimadzu visible-ultraviolet spectrophotometer (UV 2250, Fukuoka, Japan). Cell wall N content was calculated according to Onoda *et al*. [15]: 1 g of leaves were powdered in liquid nitrogen and suspended in sodium phosphate buffer (pH 7.5), the homogenate was centrifuged at 2500 g for 5 min, and the supernatant was discarded. The pellet was washed with 3% (w/v) SDS, amyloglucosidase (35 U ml^−1^, Rhizopus mold, Sigma, St Louis, MO, USA), and 0.2 M KOH, then heated and centrifuged. The pellet was then washed with distilled water and ethanol, and oven dried (75°C) for 2 days. N in the final pellet was determined using an automatic Kjeldahl apparatus (VELP Scientifica, Usmate, Italy). The fraction of leaf N allocated to cell walls (*P*_CW_) represents the ratio of cell wall N content to the total N content.

### Calculation of N Allocation in the Photosynthetic Apparatus

The fraction of leaf N allocated to Rubisco (*P*_R_,), bioenergetics (*P*_B_), and the light-harvesting components (*P*_L_) (g g^−1^) were calculated from *V*_cmax_, *J*_max_ and chlorophyll contents using the method of Niinemets and Tenhunen [7], which has been widely used in recent years [10, 59-61]:

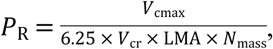

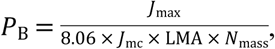

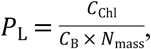

where *C*_Chl_ is the chlorophyll concentration (mmol g^−1^), *V*_cr_ is the specific activity of Rubisco (μmol CO_2_ g^−1^ Rubisco s^−1^), *J*_mc_ is the potential rate of photosynthetic electron transport (μmol electrons μmol^−1^ Cyt f s^−1^), and *C*_B_ is the ratio of leaf chlorophyll to leaf N during light-harvesting (mmol Chl (g N)^−1^). *V*_cr_, *J*_mc_, and *C*_B_ were calculated according to Niinemets and Tenhunen [7]. The fraction of leaf N allocated to the photosynthetic apparatus (*P*_P_) was calculated as the sum of *P*_R_, *P*_B_, and *P*_L_.

## Statistical Analysis

Differences between the seedling leaves were analyzed using one-way analysis of variance (ANOVA), and a post hoc test (Tukey’s test) was conducted if the differences were significant. The significance of the correlation between each pair of variables was tested with a Pearson correlation (two-tailed). All analyses were carried out using Statistical Product and Service Solutions 17.0 (SPSS17.0, Chicago, IL, USA).

## Results

### PNUE in Four Seedling Leaves

There were significant differences in PNUE between the leaves of the four seedlings (*P* < 0.001, Table 1). The PNUE in *B. alnoides* and *C. hystrix* seedling leaves were higher than those in *D. odorifera* and *E. fordii*, which was mainly attributed to their lower *N*_area_ and *N*_mass_ values. The highest PNUE in *B. alnoides* (120.54 μmol mol^−1^ s^−1^) was 2.6 times the lowest, found in *E. fordii* (45.92 μmol mol^−1^ s^−1^). However, *N*_area_ and *N*_mass_ in *B. alnoides* were 48.75% and 45.21% lower than in *E. fordii*, respectively (Table 1). There were no significant differences between *B. alnoides*, *C. hystrix*, and *D. odorifera* seedling leaves in *A*_max_*′* and the value in *E. fordii* (6.60 μmol m^−2^ s^−1^) was the smallest (Table 1). LMA of *C. hystrix* (100.13 g·m^−2^) was the highest (Table 1). *E. fordii* and *B. alnoides* seedling leaves had higher *C*_mass_ than *D. odorifera* and *C. hystrix*, but *C/N* was higher in *B. alnoides* and *C. hystrix* seedling leaves than *D. odorifera* and *E. fordii* (Table 1).

**Table 1.** Light-saturated photosynthesis (*A*_max_′), leaf N content per area (*N*_area_), leaf N content per mass (*N*_mass_), leaf C content per mass (*C*_mass_), C/N ratio, leaf mass per area (LMA), and photosynthetic-N use efficiency (PNUE) in seedling leaves of four species.

| Leaf traits | *D. odorifera* | *E. fordii* | *B. alnoides* | *C. hystrix* | *F* |
| --- | --- | --- | --- | --- | --- |
| $A_{\text{max}}'$ ($\mu\text{mol m}^{-2} \text{s}^{-1}$) | $8.04 \pm 0.46^{\text{ab}}$ | $6.60 \pm 0.50^{\text{b}}$ | $8.55 \pm 1.60^{\text{a}}$ | $8.16 \pm 0.18^{\text{ab}}$ | 3.441* |
| $N_{\text{area}}$ ($\text{g m}^{-2}$) | $2.19 \pm 0.13^{\text{a}}$ | $2.01 \pm 0.12^{\text{a}}$ | $1.03 \pm 0.25^{\text{b}}$ | $1.02 \pm 0.06^{\text{b}}$ | 36.314*** |
| $N_{\text{mass}}$ ($\text{mg g}^{-1}$) | $31.70 \pm 0.76^{\text{a}}$ | $28.09 \pm 1.49^{\text{a}}$ | $15.36 \pm 1.04^{\text{b}}$ | $10.22 \pm 0.18^{\text{c}}$ | 106.219*** |
| $C_{\text{mass}}$ ($\text{mg g}^{-1}$) | $449.50 \pm 8.86^{\text{b}}$ | $516.65 \pm 13.98^{\text{a}}$ | $493.63 \pm 5.40^{\text{a}}$ | $479.65 \pm 4.66^{\text{b}}$ | 9.713*** |
| C/N ($\text{g g}^{-1}$) | $14.24 \pm 0.48^{\text{c}}$ | $18.70 \pm 1.05^{\text{c}}$ | $32.98 \pm 2.15^{\text{b}}$ | $47.02 \pm 1.09^{\text{a}}$ | 123.492*** |
| LMA ($\text{g m}^{-2}$) | $68.97 \pm 3.90^{\text{b}}$ | $71.35 \pm 0.89^{\text{b}}$ | $67.60 \pm 5.45^{\text{b}}$ | $100.13 \pm 2.60^{\text{a}}$ | 18.272*** |
| PNUE ($\mu\text{mol mol}^{-1} \text{s}^{-1}$) | $52.64 \pm 3.78^{\text{b}}$ | $45.92 \pm 2.24^{\text{b}}$ | $120.54 \pm 5.18^{\text{a}}$ | $112.01 \pm 4.62^{\text{a}}$ | 30.833*** |

Mean values (± SD) are shown (n = 7). Different letters indicate significant differences between species (Tukey’s test, *P* < 0.05). Statistically significant *F*-ratios were denoted by \**P* < 0.05, \*\**P* < 0.01, \*\*\**P* < 0.001.

### Photosynthetic Parameters in Four Seedling Leaves

Analysis of the quantitative limitations of photosynthesis revealed that photosynthetic capacity was mainly limited by diffusional processes (*l*_s_ and *l*_m_), whereas biochemical limitations (*l*_b_) were only between 0.33% and 0.45% of the total for all studied species (Table 2).

**Table 2.** Relative stomatal (*l*_s_), mesophyll (*l*_m_) and biochemical (*l*_b_) photosynthesis limitations in four species seedling leaves.

| Leaf traits | *D. odorifera* | *E. fordii* | *B. alnoides* | *C. hystrix* | *F* |
| --- | --- | --- | --- | --- | --- |
| $I_s$ (%) | 68.09±1.35<sup>a</sup> | 58.23±1.93<sup>b</sup> | 56.84±1.67<sup>b</sup> | 56.68±2.22<sup>b</sup> | 9.001<sup>***</sup> |
| $I_m$ (%) | 31.54±1.35<sup>b</sup> | 41.39±1.95<sup>a</sup> | 42.82±1.64<sup>a</sup> | 42.87±2.22<sup>a</sup> | 8.957<sup>***</sup> |
| $I_b$ (%) | 0.36±0.05 | 0.37±0.07 | 0.33±0.04 | 0.45±0.03 | 0.964 |

Mean values (± SD) are shown (n = 7). Different letters indicate significant differences between species (Tukey’s test, *P* < 0.05). Statistically significant *F*-ratios were denoted by \**P* < 0.05, \*\**P* < 0.01, \*\*\**P* < 0.001.

Photosynthetic parameters are shown in Table 3 and Table 4. The *V*_cmax_ and *J*_max_ in *E. fordii* were higher than the other three species (Table 3) but the statistically significant values (*F*-ratios) were lower than PNUE (Table 1). Stomatal conductance (*g*_s_, 0.100 mol CO_2_ m^−2^ s^−1^) and *C*_i_ (292.88 μmol mol^−1^) in *B. alnoides* seedling leaves were higher than the other three species (Table 4). Moreover, *g*_m-Harley_ in *B. alnoides* (0.136 mol CO_2_ m^−2^ s^−1^) was higher than the other three species but *g*_m-Ethier_ (0.140 mol CO_2_ m^−2^ s^−1^) and *g*_m-Gu_ (0.160 mol CO_2_ m^−2^ s^−1^) was highest in *D. odorifera* (Table 4). The *C*_c_ in *B. alnoides* seedling leaves (all three methods) was higher than the other three species (Table 4).

**Table 3.** Maximum carboxylation rate (*V*_cmax_) and maximum electron transport rate (*J*_max_) in four species seedling leaves.

| Leaf traits | *D. odorifera* | *E. fordii* | *B. alnoides* | *C. hystrix* | *F* |
| --- | --- | --- | --- | --- | --- |
| $V_{\text{cmax}}$ (μmol m<sup>-2</sup> s<sup>-1</sup>) | 78.13±4.59<sup>b</sup> | 99.83±9.37<sup>a</sup> | 72.98±1.51<sup>b</sup> | 82.78±1.47<sup>ab</sup> | 4.786<sup>**</sup> |
| $J_{\text{max}}$ (μmol m<sup>-2</sup> s<sup>-1</sup>) | 100.71±5.80<sup>bc</sup> | 128.76±11.20<sup>a</sup> | 98.38±5.37<sup>bc</sup> | 109.27±4.05<sup>ab</sup> | 3.822<sup>*</sup> |

Mean values (± SD) are shown (n = 7). Different letters indicate significant differences between species (Tukey’s test, *P* < 0.05). Statistically significant *F*-ratios were denoted by \**P* < 0.05, \*\**P* < 0.01, \*\*\**P* < 0.001.

**Table 4.** Stomatal conductance (*g*_s_), mesophyll conductance (*g*_m_), intercellular CO_2_ concentration (*C*_i_), and CO_2_ concentration at carboxylation site (*C*_c_) in four species seedling leaves.

| Leaf traits | *D. odorifera* | *E. fordii* | *B. alnoides* | *C. hystrix* | *F* |
| --- | --- | --- | --- | --- | --- |
| $g_s$ (mol CO<sub>2</sub> m<sup>-2</sup> s<sup>-1</sup>) | 0.067±0.004<sup>bc</sup> | 0.046±0.002<sup>c</sup> | 0.100±0.013<sup>a</sup> | 0.074±0.004<sup>b</sup> | 11.106<sup>***</sup> |
| $C_i$ (μmol mol<sup>-1</sup>) | 251.54±6.44<sup>b</sup> | 235.61±6.19<sup>b</sup> | 292.88±5.94<sup>a</sup> | 256.78±5.24<sup>b</sup> | 7.348<sup>**</sup> |
| $g_{m-Harley}$ (mol CO<sub>2</sub> m<sup>-2</sup> s<sup>-1</sup>) | 0.114±0.013<sup>ab</sup> | 0.068±0.007<sup>b</sup> | 0.136±0.013<sup>a</sup> | 0.109±0.006<sup>ab</sup> | 15.391<sup>***</sup> |
| $g_{m-Ethier}$ (mol CO<sub>2</sub> m<sup>-2</sup> s<sup>-1</sup>) | 0.140±0.01<sup>a</sup> | 0.066±0.01<sup>c</sup> | 0.130±0.01<sup>ab</sup> | 0.099±0.01<sup>bc</sup> | 14.772<sup>***</sup> |
| $g_{m-Gu}$ (mol CO<sub>2</sub> m<sup>-2</sup> s<sup>-1</sup>) | 0.160±0.01<sup>a</sup> | 0.063±0.01<sup>b</sup> | 0.138±0.02<sup>a</sup> | 0.090±0.01<sup>b</sup> | 19.390<sup>***</sup> |
| $C_{c-Harley}$ (μmol mol<sup>-1</sup>) | 178.39±7.84<sup>b</sup> | 136.80±5.18<sup>c</sup> | 228.78±8.44<sup>a</sup> | 172.17±6.10<sup>b</sup> | 29.182<sup>***</sup> |
| $C_{c-Ethier}$ (μmol mol<sup>-1</sup>) | 192.99±7.11<sup>b</sup> | 133.56±7.95<sup>c</sup> | 224.91±9.13<sup>a</sup> | 173.93±3.73<sup>b</sup> | 27.639<sup>***</sup> |
| $C_{c-Gu}$ (μmol mol<sup>-1</sup>) | 200.92±6.72<sup>a</sup> | 127.42±9.56<sup>b</sup> | 225.58±12.27<sup>a</sup> | 157.86±9.56<sup>b</sup> | 20.268<sup>***</sup> |

Data of CO_2_ conductance was measured in light-saturated and atmospheric CO_2_ concentration was 380 μmol mol^−1^. Mean values (± SD) are shown (n = 7). Different letters indicate significant differences between species (Tukey’s test, *P* < 0.05). Statistically significant *F*-ratios were denoted by \**P* < 0.05, \*\**P* < 0.01, \*\*\**P* < 0.001.

### Leaf N Allocation in Four Species Seedling Leaves

There were significant differences in leaf N allocation between the four species (*P* < 0.001, Table 5). *P*_P_ was higher than *P*_CW_ in four species seedling leaves (Table 5). The *P*_P_ was 3.9 times of the *P*_CW_ in *D. odorifera*, 5.4 times in *E. fordii*, 2.0 times in *B. alnoides* and 1.6 times in *C. hystrix*. *P*_R_ > *P*_L_ > *P*_B_ in *D. odorifera*, *E. fordii*, and *B. alnoides* seedling leaves, and *P*_R_ > *P*_L_ = *P*_B_ in *C. hystrix* seedling leaves.

**Table 5.**
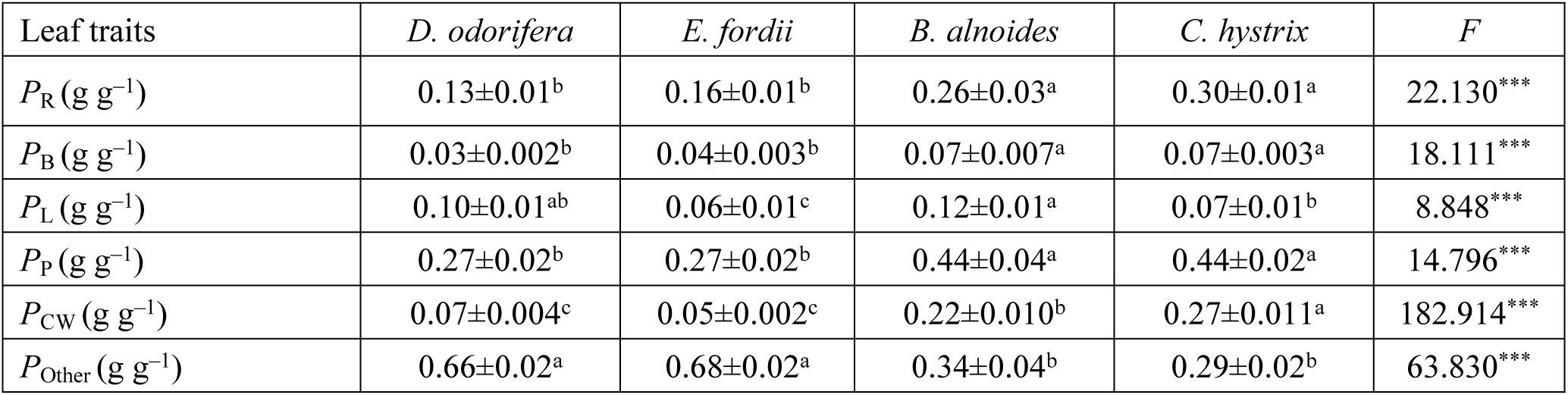
Fraction of leaf N allocated to Rubisco (*P*_R_), bioenergetics (*P*_B_), light-harvesting components (*P*_L_), photosynthetic apparatus (*P*_P_), cell wall (*P*_CW_), and other parts (1-*P*_P_-*P*_CW_, *P*_Other_) in four species seedling leaves.

Mean values (± SD) are shown (n = 7). Different letters indicate significant differences between species (Tukey’s test, *P* < 0.05). Statistically significant *F*-ratios were denoted by \**P* < 0.05, \*\**P* < 0.01, \*\*\**P* < 0.001.

The *P*_P_ in *B. alnoides* and *C. hystrix* seedling leaves (both were 0.44 g g^−1^) were higher than *D. odorifera* and *E. fordii* (both were 0.27 g g^−1^). The *P*_R_ and *P*_B_ in *B. alnoides* and *C. hystrix* seedling leaves were also higher than in *D. odorifera* and *E. fordii*. The *P*_L_ in *B. alnoides* was the highest (0.12 g g^−1^), followed by *D. odorifera* (0.10 g g^−1^), *C. hystrix* (0.07 g g^−1^), and *E. fordii* (0.06 g g^−1^).

### Relationship between PNUE and Affecting Factors

There was a positive relationship between *g*_m_ and PNUE (*P* < 0.05), in *D. odorifera, E. fordii,* and *B. alnoides*, but not in *C. hystrix* (Fig 1). Both *P*_P_, *P*_R_, and *P*_B_ had a significant positive correlation with PNUE in these species (*P* < 0.001) (Fig 2).

**Figure 1.**
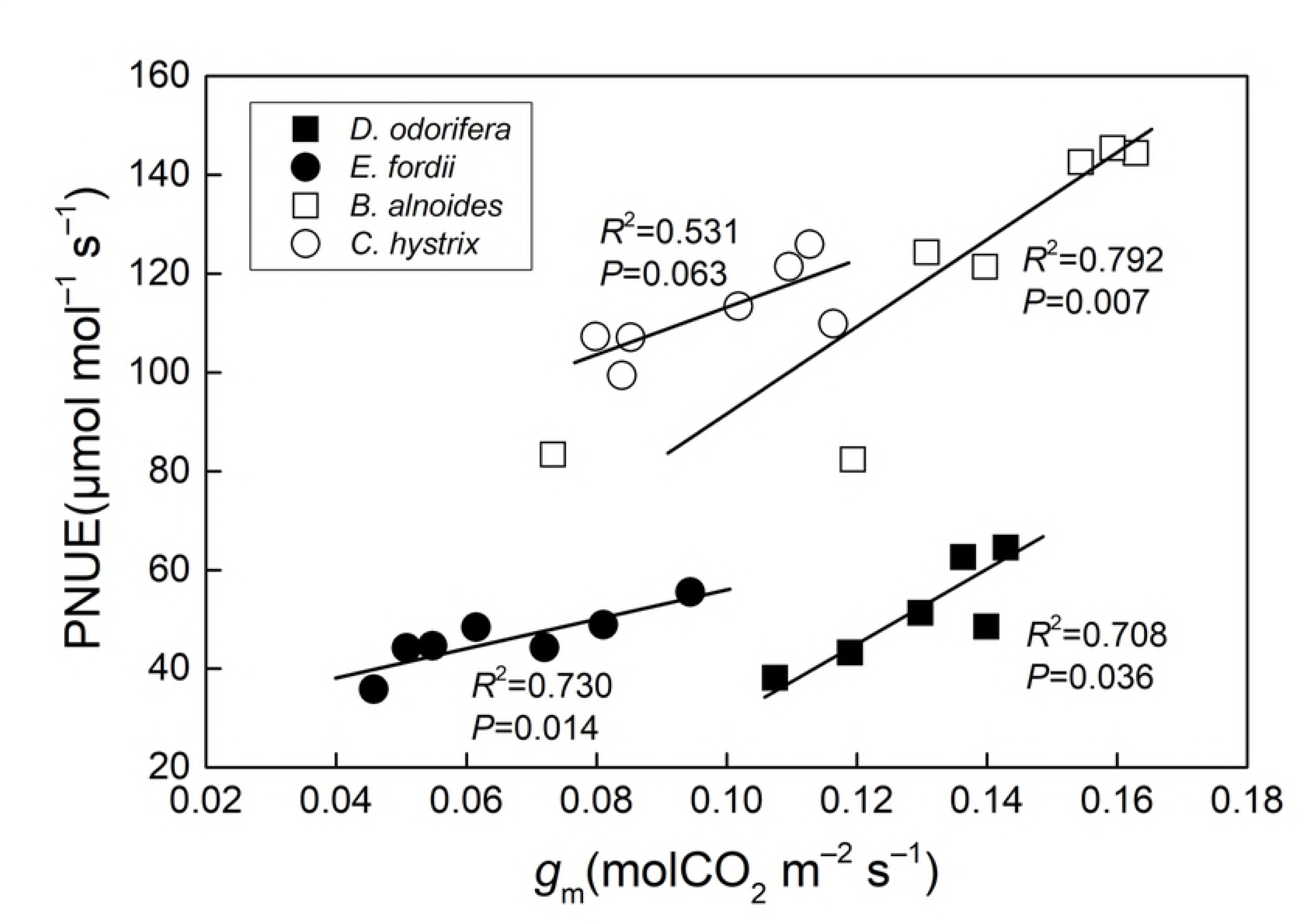
Regression analysis of mesophyll conductance (*g*_m_) with photosynthetic-N use efficiency (PNUE) in four species seedling leaves. The determination coefficient (*R*^2^) and *P*-value are also shown. The lines fitted separately for four species were significantly different (*P* < 0.05) according to the result of a one-way ANCOVA with PNUE as a dependent variable, tree species as fixed factors, and *g*_m_ as a covariate.

**Figure 2.**
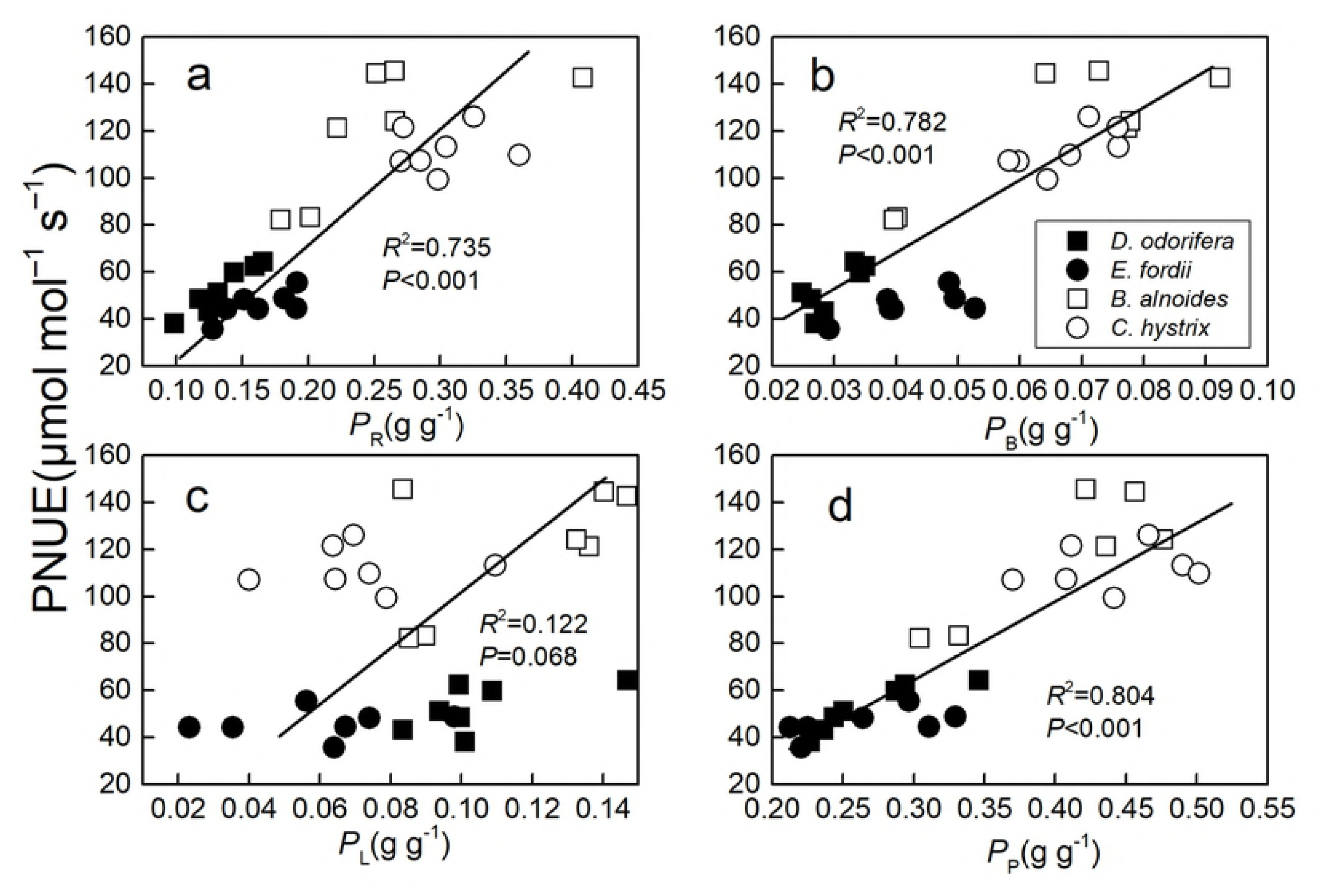
Regression analysis of the fraction of leaf N allocated to (a) Rubisco (*P*_R_), (b) bioenergetics (*P*_B_), (c) light-harvesting components (*P*_L_), and (d) the photosynthetic apparatus (*P*_P_) with photosynthetic-N use efficiency (PNUE) in four species seedling leaves. The determination coefficient (*R*^2^) and *P*-value are also shown. Only one line is fitted for four species, because there was no significant difference (*P* > 0.05) according to the result of a one-way ANCOVA with PNUE as a dependent variable, tree species as fixed factors, and *P*_R_*, P*_B_*, P*_L,_ or *P*_P_ as a covariate.

The relationship between *P*_CW_ and *P*_R_ in *B. alnoides* (*P* = 0.022) and *C. hystrix* (*P* = 0.011) seedling leaves were more significant than in *D. odorifera* (*P* = 0.409) and *E. fordii* (*P* = 0.637). Regression analysis of *P*_CW_ with *P*_R_ in *B. alnoides* seedling leaves was within the shaded zone; *C. hystrix* was on the shaded zone; *D. odorifera* and *E. fordii* were under the shaded zone (Fig 3).

**Figure 3.**
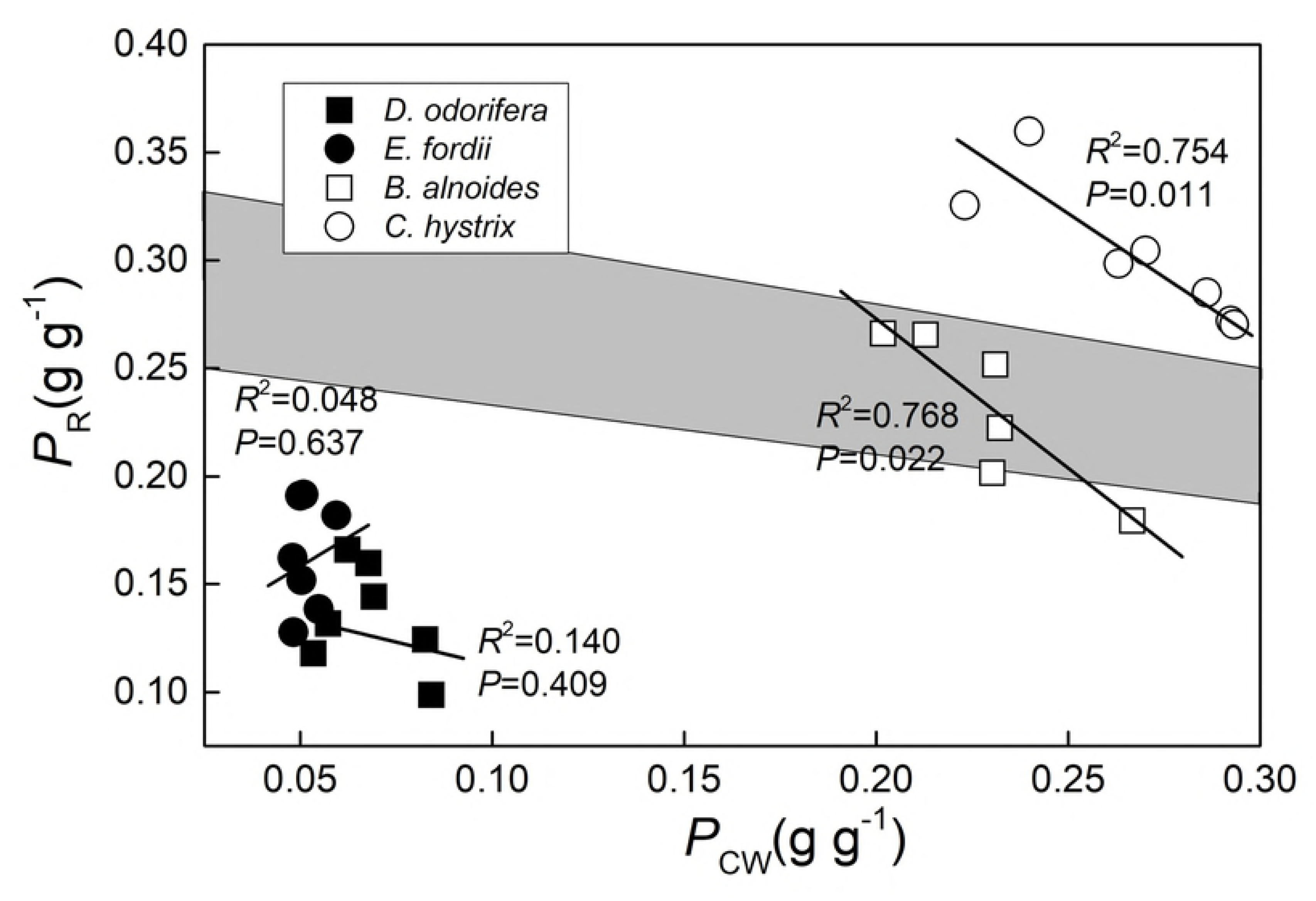
Regression analysis of the fraction of leaf N allocated to the cell wall (*P*_CW_) with leaf N allocated to Rubisco (*P*_R_) in four species seedling leaves. The determination coefficient (*R*^2^) and *P*-value are also shown. The shaded zone represents the distribution area of *P*_CW_ and *P*_R_ when a trade-off exists, for more information see Harrison *et al*. [18]. The lines fitted separately for four species were significantly different (*P* < 0.05) according to the result of a one-way ANCOVA with *P*_R_ as a dependent variable, tree species as fixed factors, and *P*_CW_ as a covariate.

## Discussion

The range of PNUE in these tree seedlings was 45.92–120.54 μmol·mol^−1^·s^−1^ (Table 1) which was close to six *Fagus sylvatica* populations (68.74–122.22 μmol·mol^−1^·s^−1^) [62] and four *Quercus* species (approximately 60–150 μmol·mol^−1^·s^−1^) [16]; lower than *P. cathayana* (171.64–213.36 μmol·mol^−1^·s^−1^) [10] and *S. alterniflora* (171.64–213.36 μmol·mol^−1^·s^−1^) [11] under different N deposition. Wright *et al*. summed up PNUE in 710 species and the range was between 10 and 500 μmol·mol^−1^·s^−1^ [63]; therefore, our results seem reasonable. Shrubs and trees usually have a low PNUE and grasses usually have high value [63]. Fast growing herbaceous species may have a PNUE higher than 200 μmol·mol^−1^·s^−1^, whereas values for evergreen woody species can be lower than 50 [3]. Our value is within the medium range.

The overall result highlights a substantial difference between N-fixing and non-N-fixing tree seedling leaves in PNUE. PNUE in *B. alnoides* and *C. hystrix* were significantly higher than that in *D. odorifera* and *E. fordii* (Table 1). The variation of PNUE may be attributable to plant evolution and natural selection [64]. Low PNUE species compensate for their low productivity with a long leaf life-span [16]; stress-tolerant species [65] and late successional species [66] usually have low PNUE values. Therefore, low PNUE in *D. odorifera* and *E. fordii* may lead to high stress-tolerance traits and increase competitiveness in poor soil [67]. Higher PNUE species such as *B. alnoides* and *C. hystrix* could grow faster [16] and have a stronger competitive ability in ecosystems with fertile soil [68].

As PNUE is the ratio of *A*_max_*′* and *N*_area_, changes of *A*_max_*′* and *N*_area_ affect PNUE. We found no significant differences in *A*_max_*′* between the four species’ seedling leaves (*F* = 3.441). Biochemical (*l*_b_) photosynthesis limitations were not significantly different between species (*F* = 0.964, Table 2) and *V*_cmax_ values were not significantly different between species (*F* ≤ 4.786, Table 3). Stomatal (*l*_s_) and mesophyll (*l*_m_) photosynthesis limitations were higher in these trees (Table 2). *D. odorifera* had higher *l*_s_ and lower *l*_m_ values than other species (Table 2) but had lower *g*_s_ and higher *g*_m_ than other species (Table 4). Therefore, limitations to CO_2_ diffusion did not change PNUE. The *g*_m_ could also influence the variation in PNUE through N allocation [22]. There was a significant positive relationship between *g*_m_ and PNUE but the effect of *g*_m_ to PNUE was not consistent between species (Fig 1). Broeckx *et al.* [25] also found this relationship in six poplar (*Populus*) genotypes, and Nha *et al.* [58] found *g*_m_ does not contribute to greater PNUE in temperate forest. The effect of *g*_m_ on PNUE was different between species.

N-fixing trees *D. odorifera* and *E. fordii* had significantly higher *N*_area_ than *B. alnoides* and *C. hystrix* (Table 1). Because *N*_area_ = *N*_mass_ × LMA, *N*_area_ may also be affected by LMA besides N content *N*_mass_. The difference of LMA between species was far lower than the difference of *N*_mass_ (Table 1). Therefore, the significantly higher *N*_mass_, caused the significantly higher *N*_area_ in *D. odorifera* and *E. fordii*. The low C/N ratio also showed high N in *D. odorifera* and *E. fordii* (Table 1). These results agreed with earlier studies [30, 31] and our study in five Fagaceae and five Leguminosae tree species [33]. However, one study reported that N-fixing trees had both higher *N*_area_ and *A*_max_*′* [29]. The relationship of *N*_area_ and *A*_max_*′* varies in patterns. The N allocation in photosynthesis was more important than the total leaf N for photosynthesis [69].

*D. odorifera* and *E. fordii* allocated a lower percentage of N to photosynthetic apparatus than *B. alnoides* and *C. hystrix*, especially to Rubisco and bioenergetics (Table 5). *P*_P_*, P*_R_, and *P*_B_ had significant positive correlations with PNUE in these trees (*R*^2^ ≥ 0.735) (Fig 2) that led to lower PNUE in N-fixing tree species (*D. odorifera* and *E. fordii*). These results agreed with previous studies [30, 31, 33, 54]. Rubisco catalyzes the limiting step for photosynthetic capacity [9]. A positive correlation between *A*_max_*′* and Rubisco has been frequently reported [12, 15]. An improved fraction of leaf N allocated to Rubisco could maximize the use of leaf N in photosynthesis. It should be noted that although there was a significant difference in N allocation proportion between N-fixing trees and non-N-fixing trees, there were smaller differences in N allocation quantity in Rubisco, bioenergetics, photosynthetic apparatus, cell wall, and other parts in the four species seedling leaves (mass and area, see S2 Table). *N*_mass_ largely affected the N allocation to the photosynthetic apparatus and *P*_CW_.

A significant negative correlation between *P*_CW_ and *P*_R_ in *B. alnoides* and *C. hystrix* (*P* < 0.05) suggested a trade-off between N allocation to Rubisco and cell walls, whereas no trade-off was detected in *D. odorifera* and *E. fordii* (Fig 3). A similar trade-off was found in *Polygonum cuspidatum* [15], *Quercus* species [16], *Mikania micrantha* and *Chromolaena odorata* [40]; but this relationship does not exist in some other trees [12]. Some researchers believe that the main influencing factors were whether leaf N could meet the needs of both cell wall N and Rubisco N [11, 18]. We used the method described by Harrison *et al.* [18] to determine whether leaf N could meet these two needs: the regression analysis of *P*_CW_ with *P*_R_ in *B. alnoides* seedling leaves was within the shaded zone (the shaded zone represents the distribution area of *P*_CW_ and *P*_R_ when a trade-off exists), *C. hystrix* was on the shaded zone which means that *B. alnoides* and *C. hystrix* had high *P*_CW_ and *P*_R_ and therefore leaf N could not meet both needs, these two factors may affect each other. We believe the high *P*_Other_ (Table 5, possibly composed of free amino acids [70] and inorganic N (NO_3_–, NH_4_+) [71]) weakens the correlation between Rubisco and cell wall N. It must be noted that *C. hystrix* showed a unique relationship between *P*_CW_ and *P*_R_ (on the shaded zone), which means higher *P*_CW_ and *P*_R_ than the results of Harrison *et al*. [18]. More trees need to be studied to determine the distribution area of *P*_CW_ and *P*_R_ when a trade-off exists.

Although both the *B. alnoides* and *C. hystrix* are non-N-fixing broadleaf plants, and had some similar functional traits, there were significant differences showed in *N*_mass_, LMA, *g*_m_, *C*_c_, and *P*_CW_ (Table 1, 4 and 5). *B. alnoides* is a deciduous broad-leaved plant and *C. hystrix* is an evergreen broad-leaf plant. In order to contribute to a longer leaf life span, evergreen broad-leaf plants should improve leaf tolerance to environmental disturbance [72,73], reflected in higher LMA [63], and *P*_CW_ [12]. Higher defensive investment could also reduce *N*_mass_ [12]. Therefore, it can be concluded that the performance of these two species is consistent with leaf economics spectrum [4, 63]. Simultaneously, if higher LMA is a result of mesophyll cell wall thickening, it will reduce *g*_m_ and *C*_c_ [74, 75], and variations in LMA are often inversely correlated with *g*_m_ and *C*_c_ [76, 77], consistent with the results of those two species.

In conclusion, four tree species seedlings have a similar photosynthetic capacity. Leaf N content and percentage of N in Rubisco and bioenergetics caused the major difference in PNUE in these tree species. Excessive storage of N in N-fixing tree species may reduce their PNUE but may be useful for future physiological processes such as reproduction [10]. Storage of N could buffer changes in other N pools such as cell wall N [15, 16, 40] (Fig 3). Evergreen tree leaves with low PNUE have multiple roles in nutrient conservation, nutrient storage, stress tolerance, herbivore deterrence, and photosynthesis [1]. We should consider that some Rubisco can also function as N storage and may not involve in photosynthesis [78, 79]. This type of Rubisco might lead to greater rates of photosynthesis under suboptimal conditions [1]. Therefore, Rubisco N calculated by the model of Farquhar *et al.* [9] might be N in activated Rubisco. Using chemical methods to extract and determine Rubisco N content could be useful [16, 80].

## Conclusions

This study indicated that PNUE was significantly lower in two N-fixing trees (*D. odorifera* and *E. fordii*) than that in two non-N-fixing trees (*B. alnoides* and *C. hystrix*). This finding was mainly attributed to lower *P*_R_ and *P*_B_. *B. alnoides* and *C. hystrix* optimized their leaf N allocation to photosynthesis. Although *g*_m_ had a significant positive correlation with PNUE in these trees, the effect of *g*_m_ on PNUE was different between species. *P*_CW_ had a significant negative correlation with *P*_R_ in *B. alnoides* and *C. hystrix* seedling leaves, but there was no significant correlation between *P*_CW_ and *P*_R_ in *D. odorifera* and *E. fordii* seedling leaves, which may indicate that *B. alnoides* and *C. hystrix* seedling leaves did not have enough N to satisfy the demand from both the cell wall and Rubisco. *B. alnoides* and *C. hystrix* with higher PNUE may have a higher competitive ability in natural ecosystems with fertile soil.

## Acknowledgments

This study was sponsored by the Fundamental Research Funds of CAF (CAFYBB2018ZA003), the National Key Research and Development Program (2016YFC0502104-02), and the projects of National Natural Science Foundation of China (31290223 and 31570240). The authors would like to thank the Experimental Center of Tropical Forestry, Chinese Academy of Forestry for providing experimental apparatus and help with measurements.

## References

1. Warren CR, Adams MA. Internal conductance does not scale with photosynthetic capacity: implications for carbon isotope discrimination and the economics of water and nitrogen use in photosynthesis. Plant Cell Environ. 2006; 29: 192–201.

2. Balotf S, Islam S, Kavoosi G, Kholdebarin B, Juhasz A, et al. How exogenous nitric oxide regulates nitrogen assimilation in wheat seedlings under different nitrogen sources and levels. Plos One. 2018; 13: e0190269.

3. Field C, Mooney HA. The photosynthesis nitrogen relationship in wild plants. In: Givnish, TJ (eds) On the Economy of Plant Form and Function: Cambridge University Press, Cambridge. 1986; pp: 25–55.

4. Wright IJ, Reich PB, Westoby M, Ackerly DD, Baruch Z, et al. The worldwide leaf economics spectrum. Nature. 2004; 428: 821–827.

5. Hikosaka K. Interspecific difference in the photosynthesis-nitrogen relationship: patterns, physiological causes, and ecological importance. J Plant Res. 2004; 117: 481–494.

6. Hikosaka K. Mechanisms underlying interspecific variation in photosynthetic capacity across wild plant species. Plant Biotechnology. 2010; 27: 223–229.

7. Niinemets Ü, Tenhunen JD. A model separating leaf structural and physiological effects on carbon gain along light gradients for the shade-tolerant species *Acer saccharum*. Plant Cell Environ. 1997; 20: 845–866.

8. Evans JR. Photosynthesis and nitrogen relationships in leaves of C3 plants. Oecologia. 1989; 78: 9–19.

9. Farquhar GD, von Caemmerer S, Berry JA. A biochemical model of photosynthetic CO_2_ assimilation in leaves of C_3_ species. Planta. 1980; 149:78-90.

10. Chen L, Dong T, Duan B. Sex-specific carbon and nitrogen partitioning under n deposition in *Populus cathayana*. Trees. 2014; 28:793-806.

11. Qing H, Cai Y, Xiao Y, Yao YH, An SQ. Leaf nitrogen partition between photosynthesis and structural defense in invasive and native tall form *Spartina alterniflora* populations: effects of nitrogen treatments. Biol Invasions. 2012; 14: 2039–2048.

12. Hikosaka K, Shigeno A. The role of Rubisco and cell walls in the interspecific variation in photosynthetic capacity. Oecologia. 2009; 160: 443–451.

13. Xu C, Rosie F, Wullschleger SD, Wilson CJ, Cai M, et al. Toward a mechanistic modeling of nitrogen limitation on vegetation dynamics. Plos One, 2012; 27: e37914.

14. Showalter AM. Structure and function of plant cell wall proteins. The Plant Cell, 1993; 5: 9–23.

15. Onoda Y, Hikosaka K, Hirose T. Allocation of nitrogen to cell walls decreases photosynthetic nitrogen-use efficiency. Funct Ecol. 2004; 18: 419–425.

16. Takashima T, Hikosaka K, Hirose T. Photosynthesis or persistence: nitrogen allocation in leaves of evergreen and deciduous *Quercus* species. Plant Cell Environ. 2004; 27: 1047–1054.

17. Onoda Y, Wright IJ, Evans JR, Hikosaka K, Kitajima K, et al. Physiological and structural tradeoffs underlying the leaf economics spectrum. New Phytol, 2017; 214: 1447–1463.

18. Harrison MT, Edwards EJ, Farquhar GD, Nicotra, AB, et al. Nitrogen in cell walls of sclerophyllous leaves accounts for little of the variation in photosynthetic nitrogen-use efficiency. Plant Cell Environ. 2009; 32: 259–270.

19. Niinemets Ü, Flexas J, Peñuelas J. Evergreens favored by higher responsiveness to increased CO_2_. Trends Ecol Evol, 2011; 26: 136–142.

20. Li Y, Gao Y, Xu X, Shen Q, Guo S. Light-saturated photosynthetic rate in high-nitrogen rice (*Oryza sativa* L.) leaves is related to chloroplastic CO2 concentration. J Exp Bot. 2009; 60: 2351–2360.

21. Xu G, Huang TF, Zhang XL, Duan BL. Significance of mesophyll conductance for photosynthetic capacity and water-use efficiency in response to alkaline stress in *Populus cathayana* seedlings. Photosynthetica. 2013; 51: 438–444.

22. Buckley TN, Warren CR. The role of mesophyll conductance in the economics of nitrogen and water use in photosynthesis. Photosynth res. 2014; 119: 77–88.

23. Flexas J, Barbour MM, Brendel O, Cabrera HM, Carriquí M, et al. Mesophyll diffusion conductance to CO2: an unappreciated central player in photosynthesis. Plant Sci. 2012; 193-194: 70–84.

24. Nakhoul NL, Davis BA, Romero MF, Boron WF. Effect of expressing the water channel aquaporin-1 on the CO2 permeability of *Xenopus oocytes*. Am J Physiol. 1998; 274: C543–C548.

25. Broeckx LS, Fichot R, Verlinden MS, Ceulemans R. Seasonal variations in photosynthesis, intrinsic water-use efficiency and stable isotope composition of poplar leaves in a short-rotation plantation. Tree Physiol. 2014; 34:701-715.

26. Harris W, Baker MJ, Williams WM. Population dynamics and competition. White Clover Wallingford, UK CAB International, 1987; 205–297.

27. Chen LY, Zhao J, Zhang RY, Wang SM. Effects of nitrogen and phosphorus fertilization on legumes in *Potentilla fruticosa* shrub in alpine meadow. Ecol Sci. 2012; 9: 512–517.

28. Reed SC. Disentangling the complexities of how legumes and their symbionts regulate plant nitrogen access and storage. New Phytol, 2017; 213, 478-480.

29. Moon M, Kang KS, Park IK, Kim T, Kim HS. Effects of leaf nitrogen allocation on the photosynthetic nitrogen-use efficiency of seedlings of three tropical species in Indonesia. J Korean Soc Appl Bi. 2015; 58:511-519.

30. Novriyanti E, Watanabe M, Makoto K, Takeda T, Hashidoko Y, et al. Photosynthetic nitrogen and water use efficiency of acacia and eucalypt seedlings as afforestation species. Photosynthetica. 2012; 50: 273–281.

31. Zhu JT, Li XY, Zhang XM, Yu Q, Lin LS. Leaf nitrogen allocation and partitioning in three groundwater-dependent herbaceous species in a hyper-arid desert region of north-western China. Aus J Bot. 2012; 60: 61–67.

32. Warren CR, Adams MA. Evergreen trees do not maximize instantaneous photosynthesis. Trends plant sci. 2004; 9: 270–274.

33. Tang JC., Cheng RM, Shi ZM, Xu GX, Liu SR, et al. Fagaceae tree species allocate higher fraction of nitrogen to photosynthetic apparatus than Leguminosae in Jianfengling tropical montane rain forest, China. Plos One, 2018: 13: e0192040.

34. Luo WY, Luo P, Liu YJ. Choice and development of the fine and valuable hardwood tree species in tropical and south subtropical regions of China. Chinese J Tropical Agriculture. 2010; 30: 15-21 (in Chinese).

35. Pang ZH. The study progress of *Betula alnoides* in China. China J Guangxi Academy Sci. 2011; 27: 243-250 (in Chinese).

36. Yang BG, Liu SL, Hao J, Pang SJ, Zhang P. Research advances on the rare tree of *Erythrophleum fordii*. Guangxi Forestry Sci. 2017; 46: 165-170 (in Chinese).

37. You Y, Huang X, Zhu H, Liu S, Liang H, et al. Positive interactions between *Pinus massoniana* and *Castanopsis hystrix* species in the uneven-aged mixed plantations can produce more ecosystem carbon in subtropical China. Forest Ecol Manag. doi:10.1016/j.foreco.2017.08.025.

38. Wang WX, Shi ZM, Luo D, Liu SR, Lu LH. Characteristics of soil microbial biomass and community composition in three types of plantations in southern subtropical area of China. Chinese J Applied Ecol. 2013; 24: 1784-1792 (in Chinese).

39. Tang JC, Shi ZM, Luo D, Liu SR. Photosynthetic nitrogen-use efficiency of *Manglietia glauca* seedling leaves under different shading levels. Acta Ecol Sinica. 2017; 37: 7493-7502 (in Chinese).

40. Zhang L, Chen X, Wen D. Interactive effects of rising CO2, and elevated nitrogen and phosphorus on nitrogen allocation in invasive weeds *Mikania, micrantha* and *Chromolaena odorata*. Biol Invasions. 2016; 18: 1391–1407.

41. Feng QH, Cheng RM, Shi ZM, Liu SR, Wang WX, et al. Response of *Rumex dentatus* foliar nitrogen and its allocation to altitudinal gradients along Balang Mountain, Sichuan, China. Chinese J Plant Ecol. 2013; 37: 591-600 (in Chinese).

42. Loreto F, Di Marco G, Tricoli D, Sharkey TD. Measurements of mesophyll conductance, photosynthetic electron transport and alternative electron sinks of field grown wheat leaves. Photosynth Res. 1994; 41: 397–403.

43. Loreto F, Tsonev T, Centritto M. The impact of blue light on leaf mesophyll conductance. J Eep Bot 2009; 112:1–8.

44. Harley PC, Loreto F, Di Marco G, Sharkey TD. Theoretical considerations when estimating the mesophyll conductance to CO2 flux by analysis of the response of photosynthesis to CO2. Plant Physiol. 1992; 98: 1429–1436.

45. Li Y, Ren BB, Ding L, Shen QR, Peng SB, Guo SW. Does chloroplast size influence photosynthetic nitrogen use efficiency?. PloS One. 2013; 8: e62036.

46. Sorrentino G, Haworth M, Wahbi S, Mahmood T, Zuomin S, Centritto M. Abscisic acid induces rapid reductions in mesophyll conductance to carbon dioxide. PloS One. 2016; 11: e0148554.

47. Momayyezi M, Guy RD. Blue light differentially represses mesophyll conductance in high vs low latitude genotypes of *Populus trichocarpa* Torr. & Gray. J Plant Physiol, 2017;213:122–128.

48. Wang X, Du T, Huang J, Peng S, Xiong D. Leaf hydraulic vulnerability triggers the decline in stomatal and mesophyll conductance during drought in rice (*Oryza sativa*). J Eep Bot. 2018; 69: 4033–4045.

49. Bernacchi CJ, Singsaas EL, Pimentel C, Portis JAR, Long SP. Improved temperature response functions for models of rubisco-limited photosynthesis. Plant Cell Environ. 2001; 24: 253–259.

50. Ethier GJ, Livingston NJ. On the need to incorporate sensitivity to CO2 transfer conductance into the farquhar-von caemmerer-berry leaf photosynthesis model. Plant Cell Environ. 2004; 27: 137–153.

51. Gu LH, Pallardy SG, Tu K, Law BE, Wullschleger SD. Reliable estimation of biochemical parameters from C3 leaf photosynthesis-intercellular carbon dioxide response curves. Plant Cell Environ. 2010; 33: 1852–1874.

52. Sharkey TD, Bernacchi CJ, Farquhar GD, Singsaas EL. Fitting photosynthetic carbon dioxide response curves for C3 leaves. Plant Cell Environ. 2007: 30: 1035–1040.

53. Loustau D, Brahim MB, Gaudillère JP, Dreyer E. Photosynthetic responses to phosphorus nutrition in two-year-old maritime pine seedlings. Tree Physiol. 1999; 19: 707–715.

54. Grassi G, Magnani F. Stomatal, mesophyll conductance and biochemical limitations to photosynthesis as affected by drought and leaf ontogeny in ash and oak trees. Plant Cell Environ. 2005; 28: 834–849.

55. Tomás M, Flexas J, Copolovici L, Galmés J, Hallik L, et al. Importance of leaf anatomy in determining mesophyll diffusion conductance to CO2 across species: quantitative limitations and scaling up by models. J Eep Bot. 2013; 64: 2269–2281.

56. Peguero-Pina JJ, Sisó S, Flexas J, Galmés J, Niinemets Ü, et al. Coordinated modifications in mesophyll conductance, photosynthetic potentials and leaf nitrogen contribute to explain the large variation in foliage net assimilation rates across *Quercus ilex* provenances. Tree Physiol. 2017; 37: 1084–1094.

57. Peguero-Pina JJ., Sisó S, Flexas J, Galmés J, García-Nogales A, et al. Cell-level anatomical characteristics explain high mesophyll conductance and photosynthetic capacity in sclerophyllous *Mediterranean oaks*. New Phytol. 2017; 214: 1–12.

58. Nha B, Hayes L, Scafaro AP, Atkin OK, Evans JR. Mesophyll conductance does not contribute to greater photosynthetic rate per unit nitrogen in temperate compared with tropical evergreen wet-forest tree leaves. New Phytol. 2018; 218: 1–13.

59. Feng YL, Wang JF, Sang WG (2007) Biomass allocation, morphology and photosynthesis of invasive and noninvasive exotic species grown at four irradiance levels. Acta Oecologica 31: 40–47.

60. Bahar NH, Ishida FY, Weerasinghe LK, Guerrieri R, O’Sullivan OS, et al. Leaf-level photosynthetic capacity in lowland Amazonian and high-elevation Andean tropical moist forests of Peru. New Phytol. 2016; doi:10.1111/nph.14079.

61. Yao HS, Zhang YL, Yi XP, Zhang XJ, Fan DY, et al. Diaheliotropic leaf movement enhances leaf photosynthetic capacity and photosynthetic use efficiency of light and photosynthetic nitrogen via optimizing nitrogen partitioning among photosynthetic components in cotton (*Gossypium hirsutum* l.). Plant Biol. 2018; 20: 213–222.

62. Sánchez-Gómez D, Robson TM, Gascó A, Gil-Pelegrín E, Aranda I. Differences in the leaf functional traits of six beech (*Fagus sylvatica*, L.) populations are reflected in their response to water limitation. Environ Exp Bot. 2013; 87: 110–119.

63. Wright IJ, Reich PB, Cornelissen JH, Falster DS, Garnier E, et al. Assessing the generality of global leaf trait relationships. New phytol. 2005; 166: 485–496.

64. Reich PB, Oleksyn J, Wright IJ. Leaf phosphorus influences the photosynthesis-nitrogen relation: a cross-biome analysis of 314 species. Oecologia. 2009; 160: 207–212.

65. Hikosaka K, Hirose T. Photosynthetic nitrogen-use efficiency in evergreen broad-leaved woody species coexisting in a warm-temperate forest. Tree Physiol. 2000; 20: 1249–1254.

66. Ellsworth DS, Reich PB. Photosynthesis and leaf nitrogen in five Amazonian tree species during early secondary succession. Ecology. 1996; 77: 581–594.

67. Kikuzawa K. A cost-benefit analysis of leaf habit and leaf longevity of trees and their geographical pattern. Am Nat. 1991; 138: 1250–1263.

68. Robinson DE, Wagner RG, Bell FW, Swanton CJ. Photosynthesis, nitrogen-use efficiency, and water-use efficiency of jack pine seedlings in competition with four boreal forest plant species. Can J Forest Res. 2001; 31: 2014–2025.

69. Qin RM, Zheng YL, Valiente-Banuet A, Callaway RM, Barclay GF, et al. The evolution of increased competitive ability, innate competitive advantages, and novel biochemical weapons act in concert for a tropical invader. New Phytol. 2013; 197: 979–988.

70. Ruan J, Haerdter R, Gerendás J. Impact of nitrogen supply on carbon/nitrogen allocation: a case study on amino acids and catechins in green tea [*Camellia sinensis* (L.) O. Kuntze] plants. Plant Biology. 2010; 12: 724–734.

71. Funk JL, Glenwinkel LA, Sack L. Differential allocation to photosynthetic and non-photosynthetic nitrogen fractions among native and invasive species. PloS one. 2013; 8: e64502.

72. Wright IJ, Cannon K. Relationships between leaf lifespan and structural defences in a low-nutrient, sclerophyll flora. Funct Ecol. 2001; 15: 351–359.

73. Onoda Y, Schieving F, Anten NP. Effects of light and nutrient availability on leaf mechanical properties of *Plantago major*: a conceptual approach. Ann Bot. 2008; 101: 727–736.

74. Warren CR. Stand aside stomata, another actor deserves centre stage: the forgotten role of the internal conductance to CO_2_ transfer. J Exp Bot. 2008; 59:1475-1487.

75. Scafaro AP, Von CS, Evans JR, Atwell BJ. Temperature response of mesophyll conductance in cultivated and wild *Oryza* species with contrasting mesophyll cell wall thickness. Plant Cell Environ. 2011;34: 1999–2008.

76. Piel C, Frak E, Le Roux X, Genty B. Effect of local irradiance on CO_2_ transfer conductance of mesophyll in walnut. J Exp Bot. 2002; 53: 2423–2430.

77. Tosens T, Niinemets U, Vislap V, Eichelmann H, Díez PC. Developmental changes in mesophyll diffusion conductance and photosynthetic capacity under different light and water availabilities in *Populus tremula*: how structure constrains function. Plant Cell Environ. 2012; 35: 839–856.

78. Stitt M., Schulze E.D. Does Rubisco control the rate of photosynthesis and plant growth? An exercise in molecular ecophysiology, Plant Cell Environ. 1994; 17: 465–487.

79. Warren CR, Adams MA. Distribution of N, Rubisco and photosynthesis in *Pinus pinaster* and acclimation to light. Plant Cell Environ. 2001; 24: 597–609.

80. Carmo-Silva E, Scales JC, Madgwick PJ, Parry MAJ. Optimizing Rubisco and its regulation for greater resource use efficiency. Plant Cell Environ. 2015; 38: 1817–1832.

